# Epigenetic state and gene expression remain stable after CRISPR/Cas-mediated chromosomal inversions

**DOI:** 10.1101/2024.10.15.618494

**Authors:** Solmaz Khosravi, Rebecca Hinrichs, Michelle Rönspies, Reza Haghi, Holger Puchta, Andreas Houben

## Abstract

In *Arabidopsis thaliana,* the chromosome arms are DNA-hypomethylated and enriched in the euchromatin-specific histone mark H3K4me3. In contrast, pericentromeric regions are DNA-hypermethylated and enriched in H3K9me2. In order to investigate how chromosomal location affects epigenetic stability and gene activity, we induced two differently-sized inversions by CRISPR/Cas and introduced heterochromatic, pericentric sequences into an euchromatic chromosomal arm. The epigenetic status of the lines was investigated by whole genome bisulfite sequencing and chromatin immunoprecipitation. Additionally, we studied the effect of the chromosomal inversions on gene expression. Our analysis revealed that both inversions affected neither the global distribution of eu- and heterochromatin-specific histone marks nor the global DNA methylation landscape. However, minor epigenetic changes were found across the entire genome. Importantly, the inverted chromosome regions and their border regions did not change their epigenetic profile. The transcription analysis of both inversion lines revealed that 0.5 - 1 % of genes were differentially expressed across the entire genome. However, the expression activity of those genes encoded by the inverted segments, and located around the cutting sites of CRISPR/Cas was less affected. Thus, gene expression levels and the epigenetic landscape remain preserved following engineered chromosomal restructuring, at least in the following generations.

## Introduction

There is a general correlation between the chromosomal position of a DNA sequence, the epigenetic state of the chromatin as well as the gene activity (de Nooijer et al., 2009). Chromosome arms are euchromatin enriched, whereas centromeric and pericentromeric regions are heterochromatic. Euchromatin, which is the decondensed fraction of chromatin, includes mostly active genes. In contrast, heterochromatin, the condensed chromatin fraction, is poor in genes and gene activity (de Nooijer et al., 2009). The formation and maintenance of the chromatin status is regulated epigenetically by DNA methylation and post-translational histone modifications. Heterochromatin is enriched in hypermethylated DNA and dimethylated histone H3K9 (H3K9me2). In contrast, euchromatin is linked with trimethylated H3K4 (H3K4me3) and less C-methylation of DNA.

Position effect variegation (PEV), discovered in the fruit fly *Drosophila melanogaster* (Gowen and Gay, 1934) and humans (Finelli et al., 2012) or telomere position effect (TPE) in budding yeast are examples of the effect of the chromosomal position on gene expression (Gottschling et al., 1990). Genes undergo differential expression in PEV because chromosomal inversions create new heterochromatin-euchromatin borders, and euchromatic genes juxtaposed to heterochromatic regions undergo heterochromatin-induced gene silencing (Hessler, 1958; Elgin and Reuter, 2013). The impact of the chromosomal position on gene expression is well-studied in case of the expression of the 45S rDNA loci in *Arabidopsis thaliana* (Mohannath et al., 2016). Other studies also suggest that changes in gene expression following the introduction of chromosomal rearrangements including inversions or translocations due to reorganization of large regulatory domains (Naseeb et al., 2016), the modification of genetic regions adjacent to their chromosome segment breakpoints (Lavington and Kern, 2017), the epigenetic environment of the translocated regions (Fournier et al., 2010) or nuclear reorganization (Fournier et al., 2010; Harewood et al., 2010). However, it is unknown whether the reported gene expression and epigenetic changes occurred immediately after the occurrence of the chromosomal rearrangements or if they were established over time in subsequent generations.

To unravel the effect of chromosomal inversions on the epigenetic state of chromatin and the activity of genes in *A. thaliana*, we employed CRISPR/Cas-assisted chromosome engineering for the generation of two differently-sized chromosomal inversions (Rönspies et al., 2022a). The inversions were first confirmed by sequencing of the inversion junctions and fluorescent *in situ* hybridization (FISH). The epigenetic state of these lines was compared to wild-type plants with the help of whole genome bisulfite sequencing (WGBS) and chromatin immune precipitation followed by sequencing (ChIP-seq) using antibodies recognizing H3K4me3 and H3K9me2 as eu- and heterochromatic histone marks, respectively. Finally, the effect of the chromosomal rearrangements on the activity of genes was analyzed. Our results showed that none of the studied inverted chromosome segments and their neighbouring regions changed in epigenetic marks and gene expression besides minor genome-wide effects, demonstrating the robustness of the epigenome and transcriptome following CRISPR/Cas-induced chromosomal restructuring, at least in the following generations.

## Results

### Targeted engineering of chromosomal inversions by CRISPR/Cas

To define whether chromosomal rearrangements influence the epigenetic state and transcriptome, we aimed to move a pericentromeric, heterochromatic region of chromosome III into an euchromatic chromosome arm environment. Two kinds of CRISPR**/**Cas engineered chromosomal rearrangements were designed for *A. thaliana* chromosome III, a 5 Mb-large paracentic and a 7.5 Mb-large pericentric inversion (Fig. 1B-C). The cut sites were chosen in a way that they were located close to the boundary of pericentromere and 178-bp satellite based on sequence information from SALK SALK 1,001 genome browser (http://signal.salk.edu/atg1001/3.0/gebrowser.php). The targeted generation of the inversions followed the protocol by (Rönspies et al., 2022a). As a first step in generating the respective inversions, suitable spacer sequences enabling high-efficiency cutting by SaCas9 had to be identified. This was particularly important for the heterochromatic pericentromeric regions of chromosome III, as heterochromatin is less accessible for Cas nucleases (Weiss et al., 2022). Several possible targets were tested by TIDE analysis (Brinkman et al., 2014) (Supplementary Table 1). In the end, three protospacers (PS) were chosen that showed a cutting efficiency of at least around 50%. The protospacer located close to the centromeric repeats (PS1) was used for the induction of both the para and pericentric inversion. SaCas9, under the control of an egg cell-specific promoter, was applied to generate heritable events inducing two simultaneous double-strand breaks on chromosome III (Steinert et al., 2015). PS1 and PS3 were used for creation of the paracentric, and PS1 and PS2 for creation of the pericentric inversion. To identify plants carrying the rearrangements, 40 T2 plant pools each were analyzed as previously described (Schmidt et al., 2020; Rönspies et al., 2022a). In the case of the paracentric inversion, four individual plants, each representing independent inversion events, out of 40 T2 pools were identified as positive. The pericentric inversion was found in one plant out of 40 T2 pools.

**Figure 1.**
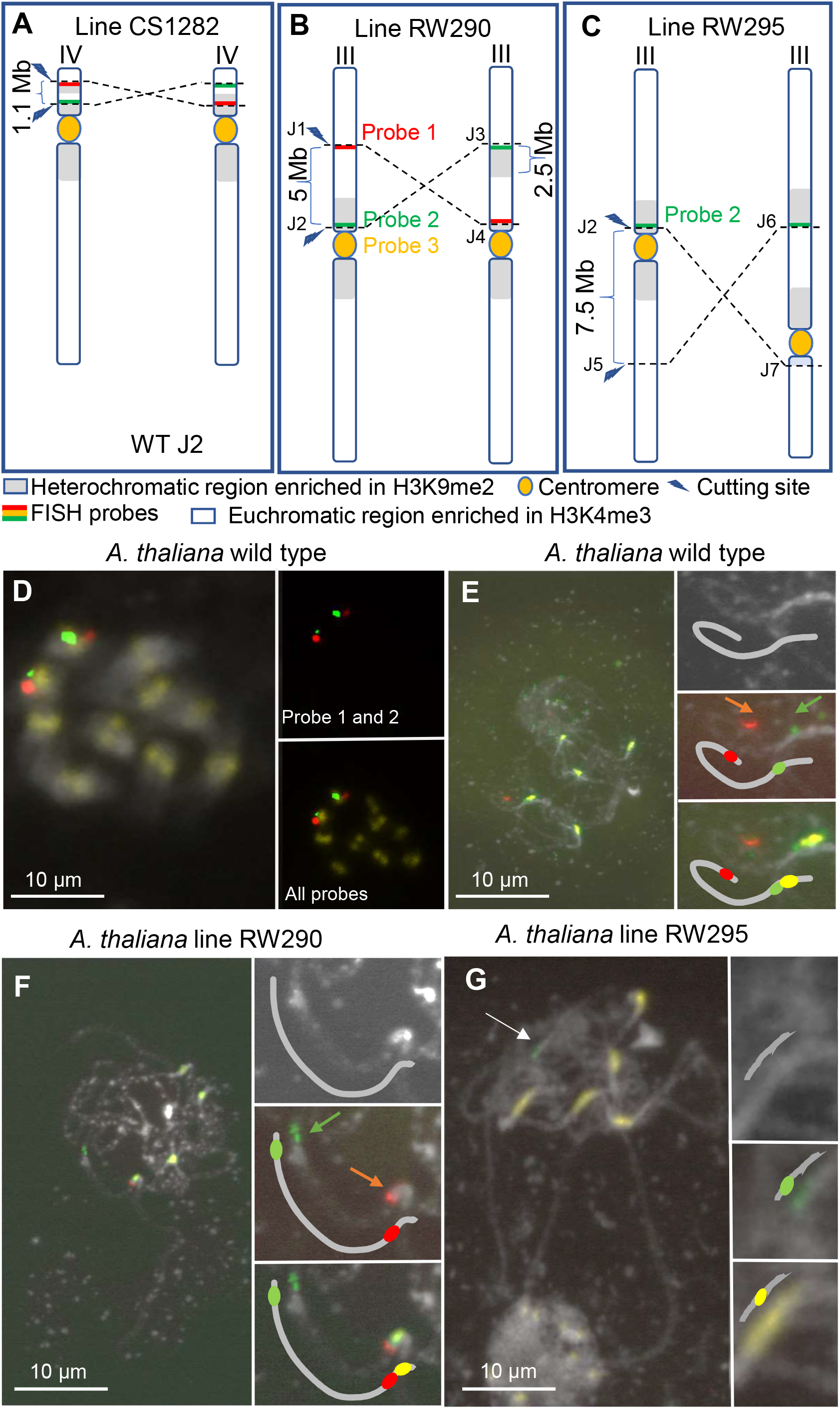
Generated and analyzed CRISPR-SaCas9-induced *A. thaliana* inversion lines. A) Line CS1282 with a re-inversion of the hk4S knob region on chromosome IV (Schmidt et al., 2020), B) line RW290 with a 5 Mb-large paracentric and C) line RW295 with 7.5 Mb-large pericentric inversions of chromosome III. Positions of the inversion breakpoints and applied FISH probes are indicated. FISH of wild-type *A. thaliana* Col 0 D) mitotic and E) pachytene chromosomes with the chromosome III- specific probes 1 (red) and 2 (green) and the centromere-specific probe 3 (yellow). F) FISH of line RW290 pachytene chromosomes with the chromosome III-specific probes 1, 2 and the centromere-specific probe. Compared to the wild-type, the green signal moved away from the centromere signal in the inversion line. Instead, the red signal moved into the vicinity of the yellow signal. G) FISH of line RW295 pachytene chromosomes with the chromosome III-specific probe 1 and the centromere-specific probe 3. Compared to the wild-type, the green signal moved away from the centromere signal in the inversion line. Chromatin was counterstained with DAPI. Insets show chromosome III further enlarged.

Next, the edited regions of the individual plants were PCR-amplified and sequenced to analyze the composition of the newly formed inversion junctions. Sequencing data revealed that one out of the four plants carrying the paracentric inversion showed a seamless ligation without any sequence loss or gain (Supplementary Fig. 1A). This plant (RW290) was chosen for further analysis. Analysis of the plant possessing the pericentric inversion revealed an insertion of one nucleotide at the break site (Supplementary Fig. 1B). To allow further experiments, both plants were propagated in the greenhouse, and the T3 seeds were sown in a germination medium without antibiotic selection. Afterwards, seedlings were genotyped via PCR using primers specific to the wild-type and inversion junctions (Supplementary Table 2). Mendelian segregation of the inversion junctions was confirmed using a chi-squared test with the critical value χ² (1; 0.95). Plants carrying the inversion in the homozygous state were cultivated until seed set.

### FISH analysis confirmed the induced chromosomal inversions

To visualize the 5 Mb-large paracentric chromosome inversion of line RW290 by FISH, oligo-painting probes (myTags libraries) were designed based on ∼100 kb-large regions downstream of the cutting site of Cas9 at position 8,592,557 (probe 1) and 100 kb upstream of the bottom cutting site of Cas9 at position 13,496,032 (probe 2) in the wild-type (Figure 1B). By analysing the order of the probes labelling the adjacent Cas9 cutting sites in combination with a centromere-specific probe (probe 3) in wild-type and line RW290 the presence of the inversion in line RW290 can be confirmed. To test the specificity of the generated FISH probes, both oligo-painting probes and the centromere-specific probe were applied to the mitotic chromosomes of wild-type Arabidopsis Col0. Accordingly, both probe 1 and 2 signals were observed on the same chromosome in the vicinity of the centromere (Figure 1D). Thus, the colocalization of probes 1 and 2 on the same chromosome proved the specificity of the generated probes that they could only label chromosome III and not any other chromosomes.

Next, to find out the order and distance of probe 1 and 2 signals relative to the centromere, pachytene chromosomes of wild-type Arabidopsis were hybridized with the same combination of FISH probes. In wild-type, the order of FISH signals generated by probes 1 and 2 was in a way that the red signal produced by probe 1 was distant from the centromere signal while the green signal generated by probe 2 was in the vicinity of the centromere signal (Fig. 1E). In line RW290, the order of the FISH signals of probe 1 and 2 was changed due to the chromosome III-specific inversion in a way that the green signal moved away from the centromere signal and the red signal moved closer to the centromere signal (Fig. 1F). Thus, the FISH analysis confirmed that line RW290 carries a CRISPR/Cas9-induced inversion on chromosome III.

To prove the presence of the 7.5 Mb-large pericentric inversion, FISH was performed with probe 1 and the centromere probe on pachytene chromosomes of line RW295 and wild-type for comparison. FISH analysis of the wild-type chromosomes showed probe 1 signals near the centromere signal (Fig. 1E). On the other hand, FISH of line RW295 with the same probes showed that the green signal from probe 1 had moved away from the centromere of chromosome III (Fig. 1G). Therefore, the FISH results proved that line RW295 indeed carries the inversion in this region.

### After induction of chromosomal inversions, the global distribution of histone marks specific to eu- and heterochromatin remains unaltered

To investigate the impact of the generated chromosome segment inversions in the earliest homozygous generation (generation T5) on the epigenetic status of the chromosomes, the distribution of post-translational histone marks typical for eu- (H3K4me3) and heterochromatin (H3K9me2) between wild-type *A. thaliana* and the inversion lines was compared. The inversion in RW290 resulted in the displacement of a 2.5 Mb-long heterochromatic region from the pericentromeric region into the euchromatic long chromosome arm. In RW295, a 7.5 Mb-long region, including the centromere of chromosome III, was inverted and translocated into the short arm of chromosome arm III, changing the submetacentric chromosome into an acrocentric chromosome. Consequently, the repositioning of eu- and heterochromatic regions in the inversion lines caused the formation of new eu /heterochromatic boundaries in chromosome III. Besides RW290 and RW295, line CS1282 (Schmidt et al., 2020) was included in this study as a control, featuring a 1.1 Mb-long inversion that moved an heterochromatic region into the heterochromatic pericentromeric region of chromosome IV (Fig. 1A).

ChIP-seq with H3K4me3- and H3K9me2-specific antibodies was performed using two-week-old seedlings of all lines to investigate the epigenetic consequences of the chromosome segment inversion. Three replicates were prepared for each genotype and antibody. The sample correlation test demonstrated a high similarity between the ChIP and input replicates. For line CS1282, only two replicates were deemed valid (Supplementary Fig. 2). To allow visual comparison of epimarks along the chromosomes, the inverted chromosome segments of all three lines are shown in an inverted orientation (Fig. 2). In other words, the chromosome segment inversions are masked. Comparison of the ChIP-seq data between inversion lines and wild-type revealed that none of the inversions affected the global distribution of H3K4me3 and H3K9me2 epimarks (Fig. 2A). In all lines, the chromosome arms were enriched in H3K4me3. At the same time, the pericentromeric regions showed a H3K9me2 enrichment. Also, at higher resolution, a comparable distribution for both epimarks was observed between the inverted chromosome segments and wild-type in the regions proximal (± 100 kb) to the breakpoints of all three inversions (Fig. 2B).

**Figure 2.**
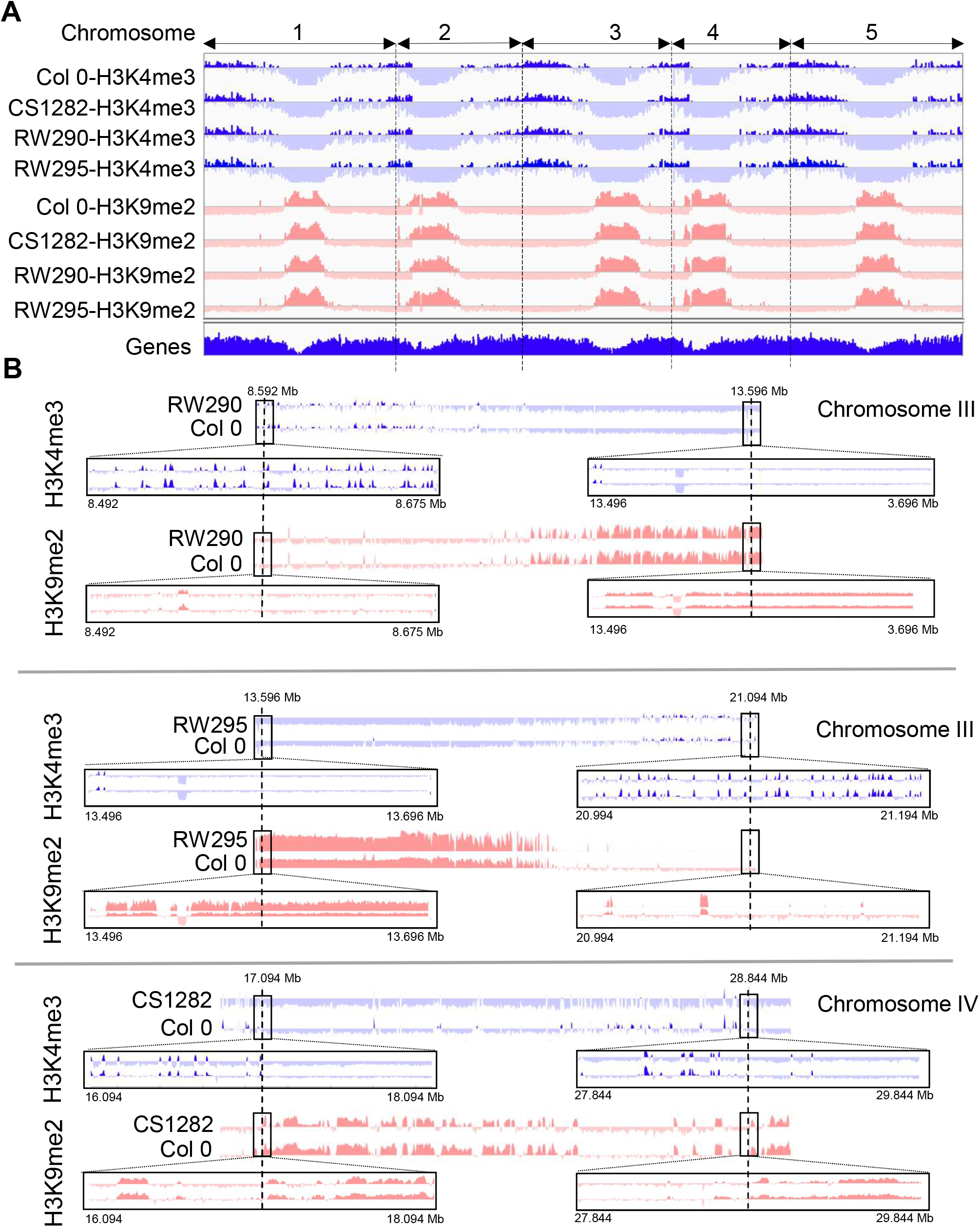
Global distribution of histone marks specific to eu- and heterochromatin remains unaltered following the induction of chromosomal inversions. A) Similar genome-wide distribution of eu- (H3K4me3) and heterochromatic (H3K9me2) histone marks between lines RW290, RW295, CS1282 and wild type *A. thaliana* Col0. B) Further resolved distribution of H3K4me3 and H3K9me2 marks within the inversion segments and proximal to the breakpoints (±100 kb). To allow visual comparison of epimarks along the chromosomes, the inverted chromosome segments of all three lines are shown in an inverted orientation. The comparisons are not in scale.

Although none of the chromosomal inversions changed the global distribution of epimarks, 29, 25 and 45 genes of lines RW290, RW295 and CS1282 changed in their histone H3K9me2 patterns compared to wild-type, respectively (Supplementary Fig. 3A, B, C). Only genes that were affected in all three replicates were considered. A slightly higher number of the altered genes was found in the case of H3K4me3. 31, 44 and 76 genes of lines RW290, RW295 and CS1282 changed in their histone H3K4me3 patterns compared to the wild-type, respectively. Further, all affected genes, reflecting an imperceptible fraction of the total number of genes, were distributed over the entire genome and not restricted to the inverted chromosome segments. Thus, except for minor exceptions, the global distribution of histone marks specific to eu- and heterochromatin remains unaltered following induction of chromosomal inversions.

### The global DNA methylome remains preserved after the induction of chromosomal inversions

To investigate the effect of the chromosome segment inversions on the DNA methylome, whole genome bisulfite sequencing (WGBS) was performed. To allow a visual comparison of methylated DNA, again, the inverted chromosome segments of all three lines are shown in an inverted orientation (Fig. 3A). The global DNA methylation profile of all three lines compared to wild-type plants was the same. Similarly, a detailed comparison of methylated CpG, CpHpG, and CpHpH sites in the inverted chromosome regions and flanking regions (± 100 kb) showed no alterations in all three lines (Fig. 3B). However, thousands of differentially methylated regions (DMRs) were found for each C context that were distributed across the entire genome. In total, 986, 729 and 901 DMRs were explicitly identified for RW290, RW295 and CS1282, respectively (Supplementary Fig.4A-B). The KEGG pathway summary of identified DMRs showed that they are mostly involved in metabolic pathways responsible for providing energy or involved in defence (Supplementary Fig. 5A-C). Thus, except for minor exceptions, the global DNA methylome remains preserved following chromosomal restructuring.

**Figure 3.**
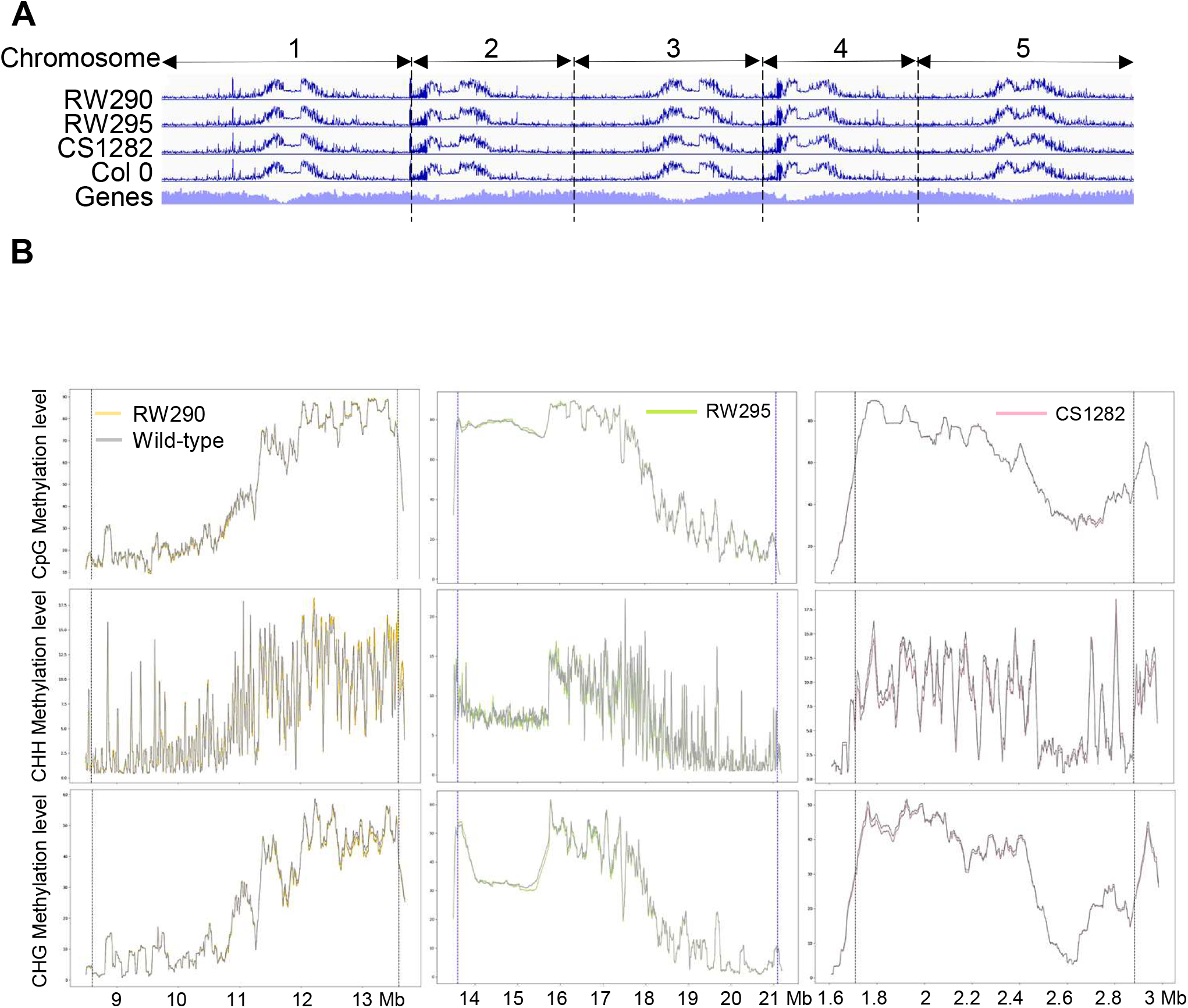
Global DNA methylome remains preserved following the induction of the chromosome segment inversions. A) Global DNA methylation pattern of all chromosomes of the lines carrying an inversion compared to the wildtype. B) Comparison of different C context methylation levels in line RW290 compared to wildtype Col0 in the area of the inversion and the ±100 kb flanking regions. The dotted blue line indicates the breakpoint positions.

### Gene expression does not change after induction of chromosome segment inversions

Finally, it was determined whether chromosome segment inversions alter the gene expression dynamics by comparative RNA-seq. The PCA analysis demonstrated a strong correlation between the replicates for each line and the distinct differences between the inversion lines and the wild-type (Supplementary Fig. 6). In each line, over 1500 differentially expressed genes (DEGs) was detected, representing 5.9 to 7.1% of the total transcriptome (Fig. 4A). Only a small number of these genes was specific to each line (Fig. 4B). In total, 0.5, 0.6 and 1.18% of DEGs were observed only in RW290, CS1282 and RW295, respectively. Therefore, the majority of DEGs were common between the three lines. The KEGG pathway enrichment analysis of DEGs revealed their involvement in metabolism or defence pathways (Supplementary Fig. 7A-C). A detailed gene activity comparison between the inverted chromosome regions and flanking regions (± 100 kb) and wild-type showed that most of the DEGs located in the inverted and flanking regions to the breakpoints not altered (Figure 4C). In total, 4, 38 and 1 DEGs were identified within the inverted segment for RW290, RW295 and CS1282, respectively. Again, most of the identified DEGs within the inverted region are involved in the regulation of metabolic pathways or defence mechanisms. Unexpectedly, the expression profile of the identified genes was not influenced by the juxtaposition of new euchromatic/heterochromatic borders (Supplementary Data Set 1). In conclusion, except for minor exceptions, the global transcriptome and epigenome remain preserved following chromosomal restructuring, at least in the following generations (Fig. 5).

**Figure 4.**
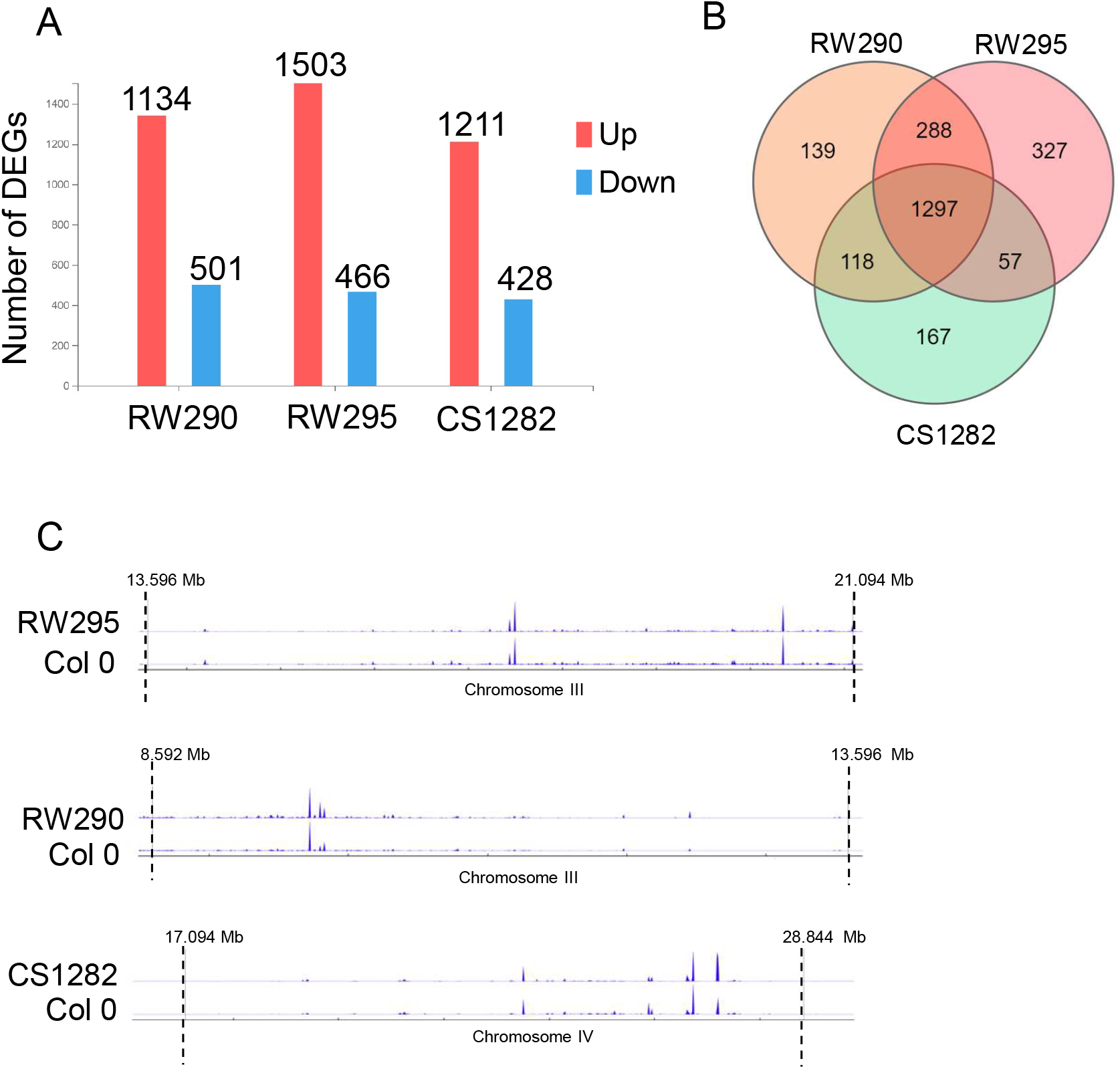
Gene expression does not change after the induction of chromosome segment inversions. A) More than 1000 differentially expressed genes were identified in each inversion line. B) From the total number of DEGs, a modest number of DEGs, specifically 139, 167 and 327 genes were recognized to be specific to lines RW290, CS1282 and RW295, respectively. 1760 DEGs were shared in all inversion lines. C) Gene expression profile of RW290, CS1282 and RW295 in the inverted and flanking regions to the break points. The expression profile of each line compared to the control was not affected by the inversion events in the inversion segments and the ±100 kb flanking regions.

**Figure 5.**
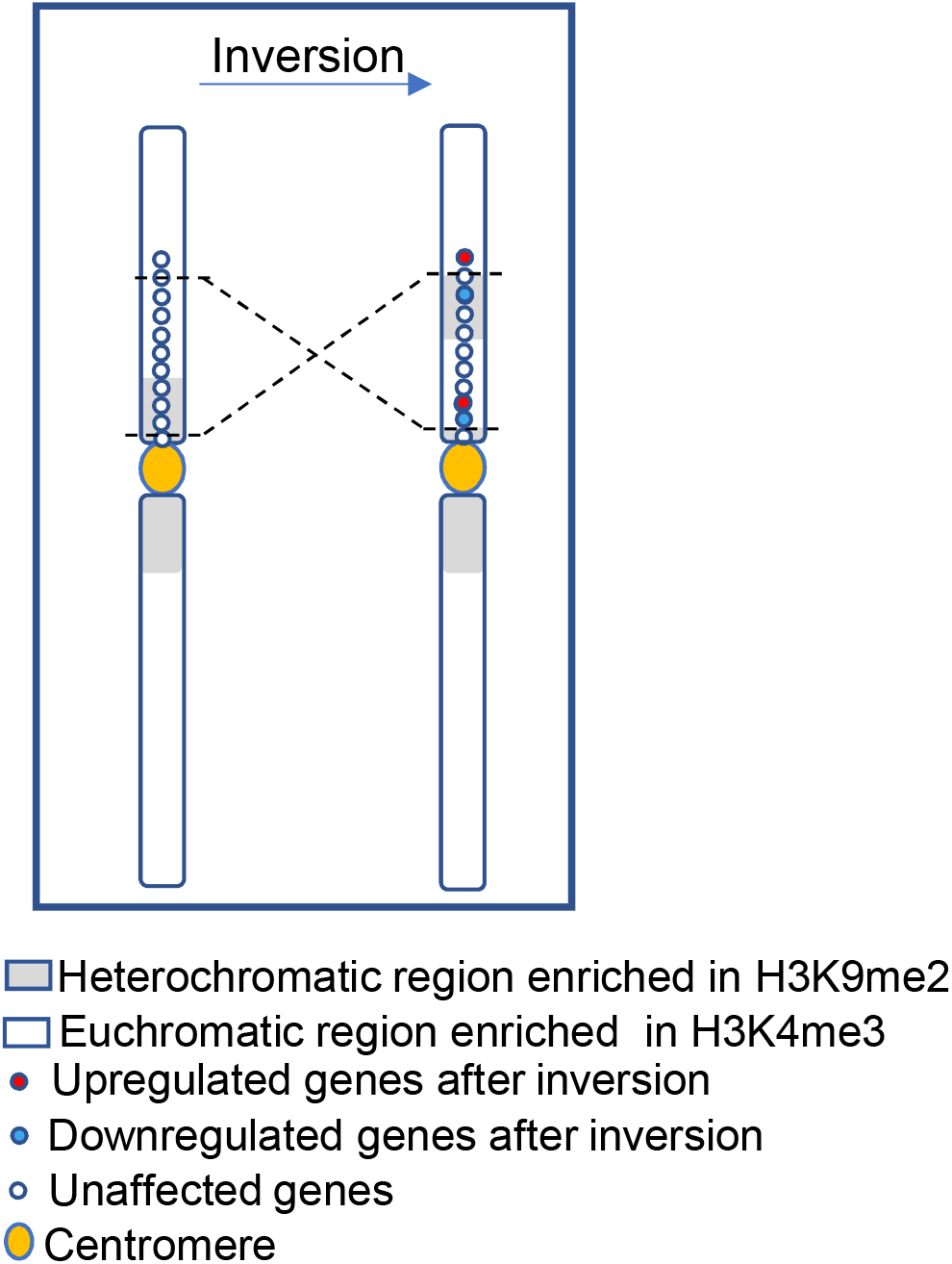
Model on the effect of a chromosome segment inversion on gene expression. Despite the formation of new eu/heterochromatic borders in the inversion lines, the genes located near the inversion borders mostly did not alter their expression due to the juxtaposition to eu-or heterochromatin except for a few genes. The activity and position of the genes are shown as red (downregulated), blue (upregulated) and white (not affected) points.

## Discussion

Chromosomal rearrangements, such as chromosome segment inversions, may affect the epigenetic landscape as well as gene expression locally and globally. Indeed, different kinds of chromosome segment inversions have been found in many cultivars of prominent crop species like rice (Zhou et al., 2023), maize (Crow et al., 2020) and barley (Jayakodi et al., 2020). So far, only historic chromosomal rearrangements that had occurred naturally could be investigated in this regard. Now that the CRISPR/Cas-based chromosome engineering technique was recently established recently, pre-defined chromosome rearrangements can be induced and their genetic and epigenetic consequences can now be analyzed directly after their occurrence (Rönspies et al., 2021). This technology is especially attractive for plant breeding, as the induction of targeted chromosomal rearrangements can be useful to manipulate genetic linkages (Puchta and Houben, 2024). By inducing reciprocal translocations between chromosomes, genetic linkages can either be broken or established (Beying et al., 2020). On the other hand, chromosomal rearrangements also play a role in the modulation of meiotic recombination as they suppress crossovers in the rearranged area during meiosis. Therefore, crossovers often present a hurdle for plant breeders since they rely on natural meiotic recombination to generate new favourable allelic combinations (Termolino et al., 2019). Thus, the possibility to reverse inversions to make recombination-dead regions of the chromosome accessible for genetic exchange again is of great value for plant breeding (Schwartz et al., 2020). Indeed, chromosome engineering could be used to revert a naturally derived 1.17 Mb inversion, called *hks4*, on chromosome 4 in *A. thaliana*. By recombination analysis, it could be shown that recombination in this previously recombination-cold area can be restored (Schmidt et al., 2020). In a second study, to determine whether targeted suppression of recombination can be achieved in a large part of the genome, almost an entire chromosome was inverted in *A. thaliana* (Rönspies et al., 2022b). The subsequent recombination analysis showed that, indeed, crossovers can be supressed in almost an entire chromosome by chromosome engineering (Rönspies et al., 2022b).

On the other hand, the application of chromosome engineering makes it possible to answer long-standing basic research questions, such as defining the role of the chromosomal position of a DNA sequence on its epigenetic stability and gene activity. To address this question, two differently-sized inversions were induced that purposely moved heterochromatic, pericentric sequences into an euchromatic chromosome arm context each. This made it possible, to test the first time whether or not the epigenetic landscape, as well as gene expression levels, remain preserved following chromosomal restructuring, at least in the following generations.

The consequences of PEV observed in *Drosophila* or TPE in budding yeast (Bao et al., 2007; Kitada et al., 2012) due to the occurrence of chromosomal rearrangements are prominent examples of the chromosome position effect on the regulation of gene expression. The underlying molecular mechanisms for the effect of chromosome position on gene expression have been attributed to several factors, including changes in the epigenetic environment of the rearranged region (Bao et al., 2007; Fournier et al., 2010; Kitada et al., 2012). In the case of PEV, heterochromatin formation depends on multiple interactions between H3K9 methyltransferases (HKMTs), heterochromatin protein 1 (HP1a) and methylation of histone H3 at lysine 9 (H3K9me2/3) (Elgin and Reuter, 2013). However, our data shows that heterochromatinization of inverted euchromatic segments juxtaposed to heterochromatic regions does not occur in the *Arabidopsis* plants that were generated in this study. The newly established eu/heterochromatic borders at the inversion points retained their wild-type epigenetic marks, including histone and methylation marks in all three analyzed inversion lines. Our finding is consistent with the effects of chromosome segment inversion observed in the hk4S genotype of *A. thaliana* and a synthetic chromosome in moss (semi-syn18L) (Fransz et al., 2016; Chen et al., 2024). However, changes in the DNA methylation profile after the occurrence of chromosomal rearrangements were reported in the case of inverted segments in human cells (Jamil et al., 2019; Carreras-Gallo et al., 2022) and *Brassica napus* hybrids (Boideau et al., 2022). Obviously, the regions selected for inversion in our experiment are not inversion prone positions as described in human cells (Jamil et al., 2019; Carreras-Gallo et al., 2022; Hazarika et al., 2022). In human cells some inversions caused diseases, although these inversions did not alter the coding sequences. Some of inversions influenced the methylation profile of the inverted segment and of the nearby borders (Jamil et al., 2019; Carreras-Gallo et al., 2022). Therefore, by changing DNA methylation the activity of genes was affected. (Carreras-Gallo et al., 2022). In addition, the effect of chromosomal rearrangements in plants has so far been studied only in the case of events that occurred many generations earlier so that chromosome inversion-independent subsequent events responsible for the changes cannot be excluded as described in *B. napus* (Jamil et al., 2019). Our findings, however, indicated that chromatin context was unaffected by inversions that occurred a few generations after the original inversion (T5). In this sense, it would be interesting to see future generations.

The perturbation of the interaction of cis- and trans-regulatory elements or the variation of genetic regions close to the inversion breakpoints are other reasons for changes in the gene activity due to the reordering of the genes’ positions in the genome (Naseeb et al., 2016; Lavington and Kern, 2017; Crow et al., 2020). In this study, the gene expression profiles showed only weak changes following the chromosomal restructuring, such as those observed in case of the hk4S inversion in *A. thaliana* (Fransz et al., 2016). In the case of the hk4S event, the inversion was induced naturally by Vandal5 transposon elements, which generated a clean split in the genes near the breakpoint (Fransz et al., 2016). In this study, the cut sites of the CRISPR/Cas system lie far beyond the regulatory regions of genes in the vicinity of the inversion points (at least for RW290) and genotyping confirmed that there is no genetic variation between the inversion and wild-type plants. This finding provides a reliable condition for focusing only on the effect of the chromosomal position on the regulation of genes. Therefore, the observed 100-300 DEGs in our lines are not likely due to the disruption of genes or their regulatory elements near the breakpoints, as they were mainly distributed throughout the genome rather than being restricted to the inversion segments or their surroundings. However, it is possible that the chromosomal rearrangements affected the 3D organization of the chromatin, and that subsequently, the expression of the underlying genes was affected slightly. In mice, it has been demonstrated that induced chromosome fusions affect the radial distribution of chromosome territories. However, these perturbations only lead to slight changes in gene expression (0.33%), with DEGs distributed globally across the genome rather than being confined to the fused chromosomes (Wang et al., 2023).

The fact that the epigenetic status of the inverted sequences was not remodelled in the generations following after the inversion event gives us the opportunity to address another important unsolved question in the future: Is the efficiency of meiotic recombination mainly determined by the position or the heterochromatic state of the respective region of the chromosome? Heterochromatic regions close to the centromere are depleted of crossovers compared to the euchromatic chromosome arms (Naish et al., 2021). By establishing the same kind of inversions as obtained in this study in a second *A. thaliana* cultivar beside Columbia, should be possible to determine by crossing and a consecutive SNP analysis whether large heterochromatic regions moved within eukaryotic chromosome arms suppress crossovers at the same level as when they were located in their original pericentric positions. This will make it possible to define whether, the chromosomal position influences crossover frequencies.

## Methods

### Generation of CRISPR/Cas9-induced *A. thaliana* inversion lines Cloning of T-DNA constructs

The Gateway-compatible plasmids pEn-Sa-Chimera and pDe-Sa-Cas9 with *Staphylococcus aureus* Cas9 under the control of an egg cell-specific promotor (pDe-Sa-Cas9 EC) were used for cloning of the transfer DNA (T-DNA) constructs (Katzen, 2007; Steinert et al., 2015; Schmidt et al., 2020). The spacer sequences were integrated into individual pEn-Sa-Chimera vectors as annealed oligonucleotides via *Bbs*I restriction digestion. The corresponding spacer sequences that are specific for both borders of the inversions are listed in Supplementary Table 1. The first guide RNA (gRNA) cassette was integrated into pDe-Sa-Cas9 EC through a classical cloning approach by *Mlu*I restriction digestion and ligation. The second gRNA cassette was transferred into the vector via a Gateway LR reaction.

### TIDE analysis

To determine the efficiency of Cas9 in inducing targeted double-strand breaks in the target regions, plants were transformed with expression constructs containing the spacer and the Cas enzyme under the control of a ubiquitin promoter. In T1 leaf material, the mutation rate was analyzed by TIDE analysis which was used as a proxy to determine the cutting efficiency (Brinkman et al., 2014). The DNA of 10 primary transformants and a Col-0 control of *A. thaliana* was extracted, and the targeted region amplified via PCR. Primers were designed in a way that they were located approximately 350 bp upstream and downstream of the predicted cleavage site. The primers are are listed in Supplementary Table 2. The reaction mixture was purified using the peqGOLD Cycle-Pure kit (VWR International, Darmstadt) and sequenced by Eurofins Genomics. Using the TIDE online tool (https://tide.nki.nl/) with default settings, the mutation rate was calculated as a proxy for the cutting efficiency in each sample. The mean value of the individual samples was calculated to determine the cutting efficiency.

### Plant cultivation and transformation

For transformation of the *A. thaliana* Col-0 plants with CRISPR/Cas expression constructs, 4-5 week-old plants were transformed via *Agrobacterium*-mediated floral dip transformation (Clough and Bent, 1998). After transformation, the plants were cultivated for 4 - 5 weeks until seed maturity. To generate sterile plant cultures, seeds were surface-sterilized with 4% sodium hypochlorite and stratified overnight at 4°C. Stratified seeds were sown on Murashige and Skoog medium (MS), 10 g/l saccharose, pH 5.7 and 7.6 g/l plant agar) containing gentamicin (0.075 g/l) and cefotaxime (0.5 g/l) to select transgenic plants. The selected transgenic plants (T1) were transferred to the greenhouse and cultivated until seed set. The harvested T2 seeds were stratified and sown on germination medium without additives, and the plates were cultivated in a growth chamber at 22°C under 16 h light/8 h dark conditions for 2 weeks.

### DNA extraction

One leaf per plant was used for DNA extraction of individual plants. To extract DNA for bulk screenings, one leaf of 40 individual plants per T2 line was collected in the same 1.5 ml reaction tube. The collected plant material was thoroughly ground with a pestle and 500 µl of extraction buffer (200 mM Tris-HCl (pH 9.0), 400 mM LiCl, 25 mM EDTA, 1% SDS, pH 9.0) was added to each tube. The blend of plant material and extraction buffer was centrifuged for 5 min at 17,000g at room temperature (RT) and the supernatant (∼400 µl) was removed and added to a new 1.5 ml tube containing 400 µl 2-propanol. The samples were mixed by inverting the tubes several times and the DNA was pelleted for 10 min at 19,500g at RT. To dry the pellets, the supernatant was discarded and the tubes with the pellets were either placed in a heating cabinet for 1.5 h at 37 °C or left to dry overnight at RT. To resuspend the DNA, 50 µl (for individual DNA extraction) or 100 µl (for bulk DNA extraction) TE buffer (10 mM Tris-HCl (pH 9.0), 1.0 mM EDTA, pH 8.0) was added to the dried pellet and agitated lightly for 15 min.

### Establishment of homozygous inversion lines

Stratified T2 seeds were sown on a germination medium and cultivated for 2 weeks in a growth chamber. Afterwards, 40 plants per T2 line were used for bulk DNA extraction. A PCR, using junction-specific primers (Supplementary Table 2), was performed on the T2 pools to screen for the presence of the inversion. If a T2 pool tested positive for the inversion, the DNA of the individual plants of this line was analyzed separately again by a PCR to identify individual plants carrying the desired restructuring. To verify the induced inversions, the junctions were sequenced by Eurofins Genomics and the software ApE (v2.0.55) was used to analyze the Sanger sequencing data by sequence alignment. Plants identified to carry the inversion were propagated in the greenhouse for 6 - 7 weeks until seed set. For genotyping in the T3 generation, PCRs were performed using specific primers for the wild-type and inversion junctions. Additionally, the T3 lines were tested for Mendelian segregation by using a chi-squared test with the critical value χ² (1; 0.95) on the genotyping results.

### Cytogenetic analysis

#### Chromosome spread preparation

Closed flower buds of ∼1 mm length were harvested and fixed in freshly prepared Carnoy’s fixative solution (3:1 v/v, ethanol: glacial acetic acid) for 48 h at RT. Chromosome spreads were prepared from fixed buds according to (Mandáková and Lysak, 2016), with the minor change of reducing the enzyme digestion time to 60 min. Prepared slides were washed with 70% ethanol for 2 min, followed with 2x SSC for 1 min. Then, they were post-fixed in 4% formaldehyde in 2x SSC for 10 min. Next, slides were washed twice in 2x SSC for 5 min and finally dehydrated in an ethanol gradient (70%, 90% and 100%, each step 2 min). The slides were air-dried for at least one hour and counterstained with DAPI (2 µg/ml in Vectashield). Finally, the slides were checked by fluorescence microscopy and the ones containing many pachytene chromosomes were selected for fluorescence *in situ* hybridization (FISH).

#### Fluorescence *in situ* hybridization (FISH)

Single-copy oligo FISH probes were designed using the Abor Biosciences’ company proprietary software (Han et al., 2015). a ∼100 kb region was selected upstream or downstream of the CRISPR/Cas9-induced break points and 45 bp long single-copy sequences were selected for the probe design. Non-overlapping target-specific oligonucleotides were synthesized as myTags libraries (Arbor Bioscience, Ann Arbor, USA). The pAL1, containing a 180 bp repeat (Martinez-Zapater et al., 1986) was labelled with Atto647N using the nick-translation labeling kit (Jena biosciences).

For FISH, slides were washed three times with 2x SSC for 5 min, dehydrated in an ethanol gradient (70%, 90% and 100%, each step 2 min), and then air dried for 1 h. In the meantime, selected myTags probes were pooled into a microtube and let to evaporate using a SpeedVac concentrator (Eppendorf). Afterwards, the probes were reconstituted in 1.5 µl of ddH_2_O. Per slide, 1400 ng per myTags probe was used. Before adding the myTags probe, 75 ng centromere-specific probe was added to 18.5 µl of hybridization mixture (50% formamide; 10% dextran sulfate; 10% salmon sperm DNA; 2×SSC) per slide and, denatured at 95 °C for 10 min and then placed on ice for 5 min. Next, the reconstituted myTags probes were added to the mixture. The entire volume of the prepared probe mixture (myTags and centromere probes) was added to the slide and covered with a coverslip. Slides were incubated for 20 min at 37°C in a wet chamber and then denatured on a hot plate (70°C) for 3 min. Finally, slides were sealed with rubber cement, placed into a wet chamber and hybridized for 48 h at 37 °C. To remove the coverslips, the slides were washed in 2x SSC at RT for 5 min. Post-hybridization washing was carried out by washing in 2x SSC at 42°C for 20 min under shaking conditions and finally washed in 2× SSC at RT for 5 min in darkness. At the end, the slides were dehydrated in an ethanol series (70%, 90% and 96%, each step 2 min), air-dried in the dark and counterstained with 8 µl DAPI (2 µg/ml in Vectashield). Micrographs were captured using an epifluorescence microscope (Olympus BX61) equipped with a cooled charge-coupled device (CCD) camera (Orca ER; Hamamatsu) and pseudo-coloured by the Adobe Photoshop 6.0 software.

#### Plant growth condition for RNA-seq and epigenome analysis

For comparative RNA-seq and epigenome analysis, seeds of *A. thaliana* wild-type (Col 0), line CS1282 (Schmidt et al., 2020), (Fig 1A), and the newly generated inversion lines were surface sterilized and cultured on MS medium. The seeds were stratified at 4°C for one night and were afterwards cultivated in a growth chamber at 22°C under 16/8 h light/dark conditions for two weeks. Two-week-old seedlings were harvested and immediately flash-frozen in liquid nitrogen. Three replicates were collected per line.

#### RNA-seq analysis

Total RNA was extracted from 100 mg of ground tissue following the protocol of the Quick-RNA Miniprep kit (Zymo Research). The integrity of the RNA samples was assessed using the RNA Integrity Number (RIN). The RNA samples were sequenced using the DNBseq PE150 platform by BGI (Hong Kong, China). A total of 50 million reads was generated for each sample. The bioinformatics analyses of data were conducted using the Dr Tom network platform provided by BGI (http://report.bgi.com). Quality control measures were applied, and adapter sequences were trimmed using SOAPnuke. Subsequently, the clean data were aligned to the reference genome of *A. thaliana* (Naish et al., 2021) using HISAT2 (Kim et al., 2015). Differential expression analysis was performed between the inversion lines and the wild-type using the DESeq2 package (Love et al., 2014) with a significance threshold of Q-value ≤ 0.01 and an absolute log2 fold change (|log2FC|) ≥ 2. Furthermore, the annotated genes were subjected to KEGG pathway enrichment analysis to elucidate their functional significance (Kanehisa et al., 2008). Plots of inverted segments and their flanking by regions showing the profile of DEGs were done with pyGenomeTracks (Lopez et al., 2021).

#### Chromatin immunoprecipitation followed by sequencing (ChIP-seq) and analysis

To extract chromatin, 1 g of ground tissues from two-week old seedlings was utilized. The ChIP followed the protocol described by (Kuo et al., 2023), with an increased fixation time to 25 minutes and the use of a total of 28 cycles for sonication. Antibodies targeting histone H3K4me3 (ab8580, Abcam) and H3K9me2 (ab1220, Abcam) were used to enrich eu- and heterochromatin (1 µl of antibody for each 100 µl of chromatin), respectively. The concentration of extracted chromatin was quantified using the QubitTM dsDNA HS Assay kit (Invitrogen, USA). For the preparation of ChIP-seq libraries, 3 ng of chromatin per sample was used, following the instructions provided by NEBNext Ultra II DNA Library Prep Kit (NEB, E7645). Subsequently, the libraries were sequenced using DNBseq PE150 by BGI (Hong Kong, China), generating 25 million reads for each library. Three replicates were prepared for each ChIP experiment.

For bioinformatic analysis, the tools available in the Galaxy portal (https://galaxy.ipk-gatersleben.de) were utilized as described by (Freeberg M. et al.). First, quality control and adapter trimming of ChIP-seq and input reads were performed with FastQC (Version 0.11.8) and Trimmomatic, (Version 0.38), respectively. Afterwards, paired-end reads (2x 150 bp) were aligned to the *A. thaliana* genome (Naish et al., 2021) using Bowtie2 with default parameters (Langmead and Salzberg, 2012). MultiBamSummary (Version 3.3.0.0.0) was used based on Pearson’s correlation coefficient to assess the similarity between the replicates of each group. Peak calling was executed using MACS2 (Version 2.1.1.20160309.6) (Zhang et al., 2008) with the following parameters: Effective genome size: 119482012, Lower mfold: 10, upper mfold: 30, minimum FDR: 0.05, Composite broad regions: broad, Duplicate tags at the exact same location: 1 . Peaks associated with H3K4me3 were analyzed as narrow peaks by adjusting composite broad regions to: no broad. To identify differentially marked genes with H3K4me3 and H3K9me2, the information from two replications was analyzed by Diffbind (Stark and Brown, 2012). Genes with p-value < 0.05 and log_2_ (FC)> 2 were recognized as differentially methylated H3K4me3 and H3K9me2. The GO and KEGG analysis of genes associated with identified unique peaks in the inversion lines was conducted using the Database for Annotation, Visualization and Integrated Discovery (DAVID) (Huang da et al., 2009; Sherman et al., 2022). Normalized coverage BIGWIG files, representing the normalized read coverage across the genome, were generated using BamCompare (Version 3.3.0.0.0) by calculating the average log2-ratio of read counts from ChIP over input (Ramírez et al., 2016). The generated normalized BIGWIG files were visualized by IGV and pyGenomeTracks to illustrate the distribution of mapped reads across the genome (Robinson et al., 2011; Lopez-Delisle et al., 2020). The pathway enrichment bubble plots and KEGG summery plots were generated using SRplot platform (https://www.bioinformatics.com.cn/en) to illustrate the enrichment of specific biological pathways in the analyzed data (Tang et al., 2023).

#### DNA methylation analysis

DNA extraction was performed using the DNeasy® Plant Mini Kit. (Qiagen). Three replicates were included for each line. The concentration of DNA was quantified using the Qubit dsDNA broad-range kit (Invitogen). To assess DNA methylation patterns, the samples were sent to BGI (Hong Kong, China), for whole-genome bisulfite sequencing. Prior to analysis, the raw sequencing data underwent quality assessment using FastQC and subsequent trimming with Trim-Galore. The reads were aligned to the reference using the Bismark pipeline (Krueger and Andrews, 2011) followed by methylation calling using methylpy with specific parameters: min-num-dms 10, min-cov 5, sig-cutoff 0.001, dmr-max-dist 200. Visualization of the data was facilitated using IGV and pyGenomeTracks (Robinson et al., 2011; Lopez-Delisle et al., 2020). Functional annotation of differentially methylated genes was performed using DAVID (Sherman et al. 2022; Huang et al. 2009).

## DATA AVAILABILITY

The authors declare that the data supporting the findings are available within the paper and its Supplementary Information, or are available from the corresponding author upon reasonable request.

## ACKNOWLEDGMENTS

We thank Katrin Kumke, Oda Weiss, Pia Kunz, Carolin Brechtel and Kristina Riedinger for their technical assistance in performing experiments. This research was funded by the BMBF project EPICHROM to HP and AH.

## Contributions

AH, SK and HP designed research; RHi generated CRISRP-Cas engineered chromosome segment inversion; SK performed FISH, ChIP-seq and analyzed genomic data; RHa analysed genomic data; and all authors wrote the paper.

---

**Supp. Figure 1.** Molecular nature of the wild-type and inversion junctions of the inversion lines. A) Molecular nature of the wild-type (WT J1 and J2) and inversion junctions (RW290 J3 and J4) of the paracentric inversion. The first two lines show the natural wild-type conformation and the last two lines show the nucleotide composition of the inversion junctions in line RW290 as determined by Sanger sequencing. The protospacer adjacent motif (PAM) of the 5′-sequence is highlighted in pink and the corresponding spacer sequence is highlighted in blue. The PAM of the 3′-sequence is highlighted in red and the corresponding spacer sequence is highlighted in green. B) Molecular nature of the wild-type (WT J2 and J5) and inversion junctions (RW295 J6 and J7) of the pericentric inversion. The first two lines show the natural wild-type conformation and the last two lines show the nucleotide composition of the inversion junctions in line RW295 as determined by Sanger sequencing. The PAM of the 5′- sequence is highlighted in red and the corresponding spacer sequence is green in blue. The PAM of the 3′-sequence is highlighted in pink and the corresponding spacer sequence is highlighted in yellow. The red letters represent the insertion of one nucleotide at the break site. Since PS1 was used to generate both inversions, the WT J2 sequence is identical in A) and B).

**Supp. Figure 2.** Sample correlation test between replicates of ChIP and Input samples.

**Supp. Figure 3.** Number of genes with differentially K4 and K9 methylated histone marks demonstrated for three replicates of ChIP-seq in line A) RW290, B) CS1282 compared to wild-type.

**Supp. Figure 4.** RW295, RW290 and CS1282 inversion line-specific genome-wide distributed DMRs. A) Comparison of the number of shared and inversion line-specific DMRs identified in each line. B) Comparison of methylation level in different C contexts between WT and inversion lines. Compared to WT, the inversion lines show a slight reduction in methylation levels of CG, CHG and CHH.

**Supp. Figure 5.** KEGG pathway summary of identified DMRs in A) line RW290 and B) Rw295 and C) CS1282. The identified genes are mostly involved in the regulation of metabolism pathways.

**Supp. Figure 6.** PCA test comparing the transcriptome of *Arabidopsis* lines RW295 and RW290 with the wild-type.

**Supp. Figure 7.** KEEG pathway enrichment histogram of recognized DEGs for lines A) RW290, B) RW295 and C) CS1282.

**Supplementary Table 1.**
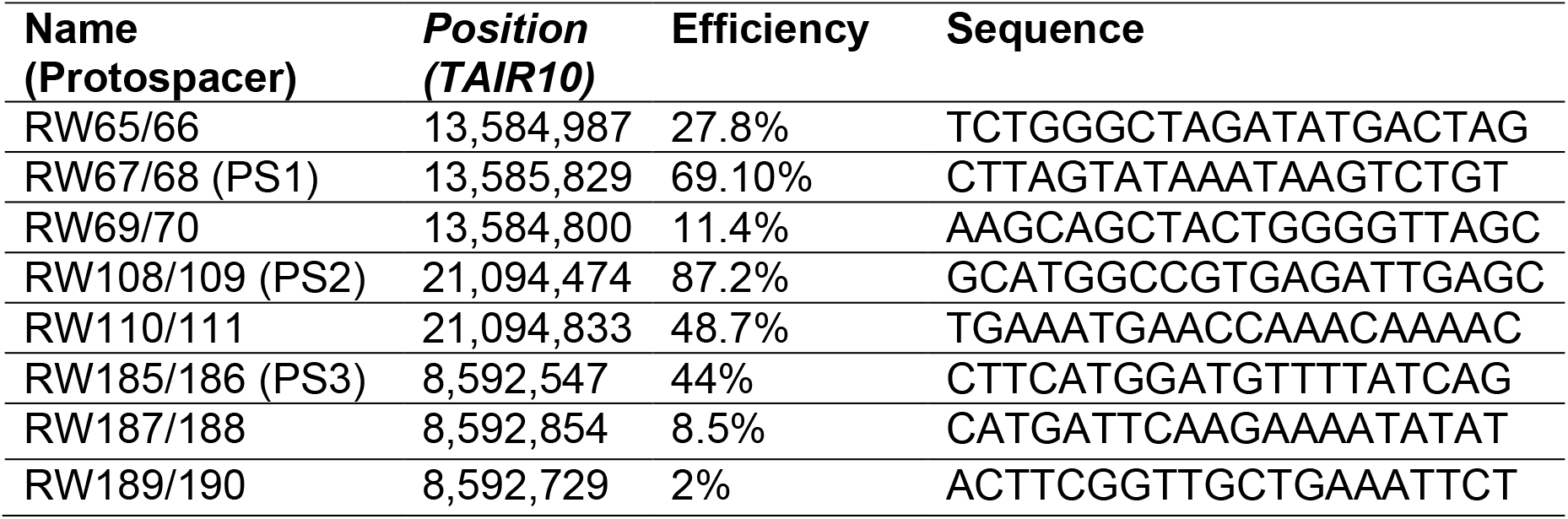
List of the protospacers tested for the establishment of both inversions. Shown here are the name, the positions of the protospacer on chromosome 3 (TAIR10 as reference genome), the cut efficiency and sequence of the protospacers. Highlighted green are the protospacers with the highest cutting efficiency, which were used to generate the here introduced inversions. The protospacer RW67/68 (PS1) was used in combination with RW108/109 (PS2) to generate the pericentric inversion and in combination with RW185/186 (PS3) to establish the pericentromeric inversion.

**Supplementary Table 2.**
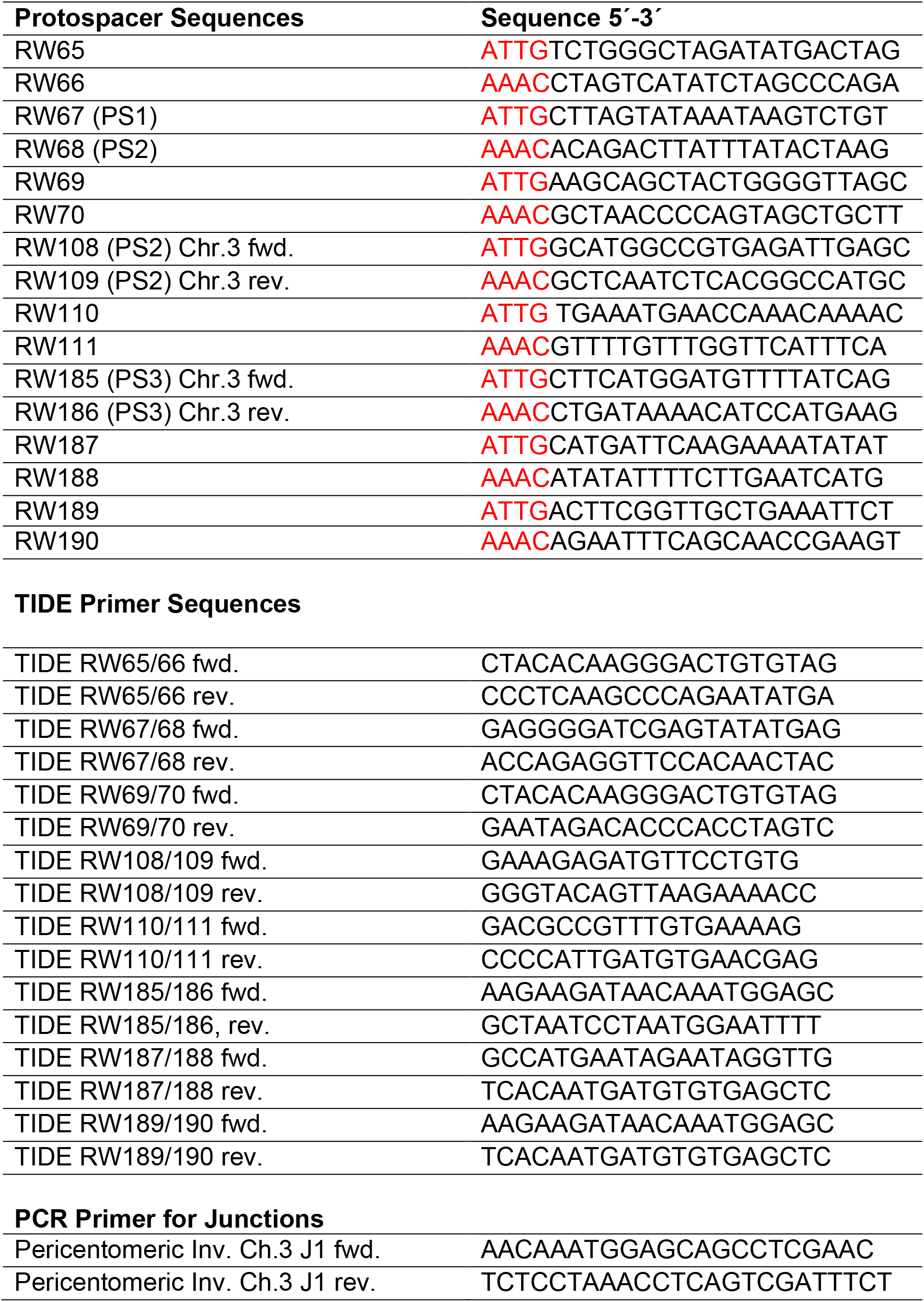
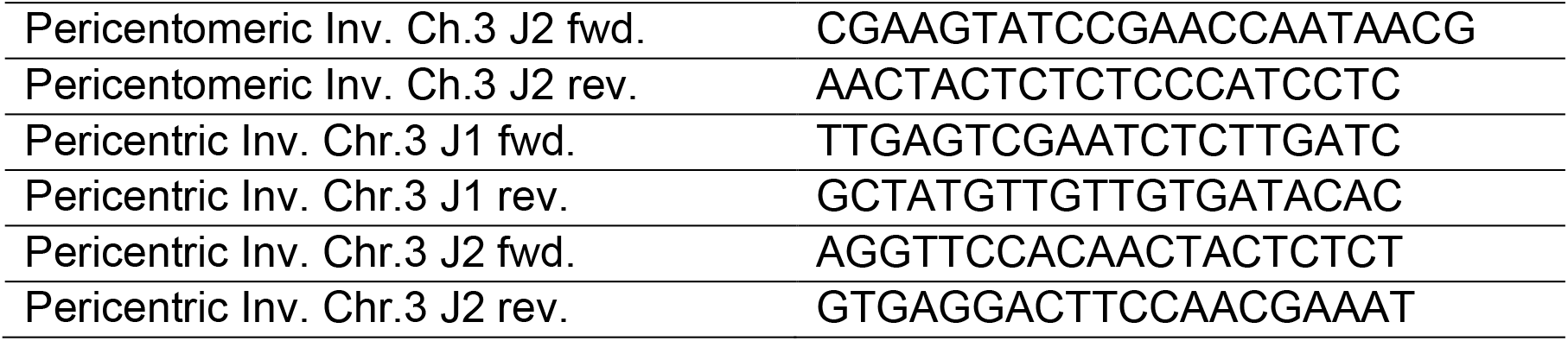
The sequences which were used as protospacer, TIDE primer and PCR primers for junctions.

## References

1. Bao, X., Deng, H., Johansen, J., Girton, J., and Johansen, K.M. (2007). Loss-of-function alleles of the JIL-1 histone H3S10 kinase enhance position-effect variegation at pericentric sites in Drosophila heterochromatin. Genetics **176**, 1355–1358.

2. Beying, N., Schmidt, C., Pacher, M., Houben, A., and Puchta, H. (2020). CRISPR–Cas9-mediated induction of heritable chromosomal translocations in Arabidopsis. Nature Plants **6**, 638–645.

3. Boideau, F., Richard, G., Coriton, O., Huteau, V., Belser, C., Deniot, G., Eber, F., Falentin, C., Ferreira de Carvalho, J., Gilet, M., Lodé-Taburel, M., Maillet, L., Morice, J., Trotoux, G., Aury, J.-M., Chèvre, A.-M., and Rousseau-Gueutin, M. (2022). Epigenomic and structural events preclude recombination in Brassica napus. New Phytologist **234**, 545–559.

4. Brinkman, E.K., Chen, T., Amendola, M., and van Steensel, B. (2014). Easy quantitative assessment of genome editing by sequence trace decomposition. Nucleic Acids Research **42**, e168–e168.

5. Carreras-Gallo, N., Cáceres, A., Balagué-Dobón, L., Ruiz-Arenas, C., Andrusaityte, S., Carracedo, Á., Casas, M., Chatzi, L., Grazuleviciene, R., Gutzkow, K.B., Lepeule, J., Maitre, L., Nieuwenhuijsen, M., Slama, R., Stratakis, N., Thomsen, C., Urquiza, J., Wright, J., Yang, T., Escaramís, G., Bustamante, M., Vrijheid, M., Pérez-Jurado, L.A., and González, J.R. (2022). The early-life exposome modulates the effect of polymorphic inversions on DNA methylation. Communications Biology **5**, 455.

6. Chen, L.-G., Lan, T., Zhang, S., Zhao, M., Luo, G., Gao, Y., Zhang, Y., Du, Q., Lu, H., Li, B., Jiao, B., Hu, Z., Ma, Y., Zhao, Q., Wang, Y., Qian, W., Dai, J., and Jiao, Y. (2024). A designer synthetic chromosome fragment functions in moss. Nature Plants **10**, 228–239.

7. Clough, S.J., and Bent, A.F. (1998). Floral dip: A simplified method for Agrobacterium-mediated transformation of Arabidopsis thaliana. The Plant Journal **16**, 735–743.

8. Crow, T., Ta, J., Nojoomi, S., Aguilar-Rangel, M.R., Torres Rodríguez, J.V., Gates, D., Rellán-Álvarez, R., Sawers, R., and Runcie, D. (2020). Gene regulatory effects of a large chromosomal inversion in highland maize. PLoS Genet **16**, e1009213.

9. de Nooijer, S., Wellink, J., Mulder, B., and Bisseling, T. (2009). Non-specific interactions are sufficient to explain the position of heterochromatic chromocenters and nucleoli in interphase nuclei. Nucleic Acids Research **37**, 3558–3568.

10. Elgin, S.C., and Reuter, G. (2013). Position-effect variegation, heterochromatin formation, and gene silencing in Drosophila. Cold Spring Harb Perspect Biol **5**, a017780.

11. Finelli, P., Sirchia, S.M., Masciadri, M., Crippa, M., Recalcati, M.P., Rusconi, D., Giardino, D., Monti, L., Cogliati, F., Faravelli, F., Natacci, F., Zoccante, L., Bernardina, B.D., Russo, S., and Larizza, L. (2012). Juxtaposition of heterochromatic and euchromatic regions by chromosomal translocation mediates a heterochromatic long-range position effect associated with a severe neurological phenotype. Molecular Cytogenetics **5**, 16.

12. Fournier, A., McLeer-Florin, A., Lefebvre, C., Duley, S., Barki, L., Ribeyron, J., Alboukadel, K., Hamaidia, S., Granjon, A., Gressin, R., Lajmanovich, A., Bonnefoix, T., Chauvelier, S., Debernardi, A., Rousseaux, S., de Fraipont, F., Figeac, M., Kerckaert, J.P., De Vos, J., Usson, Y., Delaval, K., Grichine, A., Vourc’h, C., Khochbin, S., Feil, R., Leroux, D., and Callanan, M.B. (2010). 1q12 chromosome translocations form aberrant heterochromatic foci associated with changes in nuclear architecture and gene expression in B cell lymphoma. EMBO Molecular Medicine **2**, 159–171.

13. Fransz, P., Linc, G., Lee, C.R., Aflitos, S.A., Lasky, J.R., Toomajian, C., Ali, H., Peters, J., van Dam, P., Ji, X., Kuzak, M., Gerats, T., Schubert, I., Schneeberger, K., Colot, V., Martienssen, R., Koornneef, M., Nordborg, M., Juenger, T.E., de Jong, H., and Schranz, M.E. (2016). Molecular, genetic and evolutionary analysis of a paracentric inversion in Arabidopsis thaliana. Plant J **88**, 159–178.

14. Freeberg M., Heydarian M., Bhardwaj F., Wolff J., and Erxleben A. Identification of the binding sites of the T-cell acute lymphocytic leukemia protein 1 (TAL1) (Galaxy Training Materials). https://training.galaxyproject.org/training-material/topics/epigenetics/tutorials/tal1-binding-site-identification/tutorial.html

15. Gottschling, D.E., Aparicio, O.M., Billington, B.L., and Zakian, V.A. (1990). Position effect at S. cerevisiae telomeres: Reversible repression of Pol II transcription. Cell **63**, 751–762.

16. Gowen, J.W., and Gay, E.H. (1934). Chromosome Constitution and Behavior in Eversporting and Mottling in Drosophila Melanogaster. Genetics **19**, 189–208.

17. Han, Y., Zhang, T., Thammapichai, P., Weng, Y., and Jiang, J. (2015). Chromosome-Specific Painting in Cucumis Species Using Bulked Oligonucleotides. Genetics **200**, 771–779.

18. Harewood, L., Schütz, F., Boyle, S., Perry, P., Delorenzi, M., Bickmore, W.A., and Reymond, A. (2010). The effect of translocation-induced nuclear reorganization on gene expression. Genome Res **20**, 554–564.

19. Hazarika, R.R., Serra, M., Zhang, Z., Zhang, Y., Schmitz, R.J., and Johannes, F. (2022). Molecular properties of epimutation hotspots. Nat Plants **8**, 146–156.

20. Hessler, A.Y. (1958). V-Type Position Effects at the Light Locus in Drosophila Melanogaster. Genetics **43**, 395–403.

21. Huang da, W., Sherman, B.T., and Lempicki, R.A. (2009). Systematic and integrative analysis of large gene lists using DAVID bioinformatics resources. Nat Protoc **4**, 44–57.

22. Jamil, M.A., Sharma, A., Nuesgen, N., Pezeshkpoor, B., Heimbach, A., Pavlova, A., Oldenburg, J., and El-Maarri, O. (2019). F8 Inversions at Xq28 Causing Hemophilia A Are Associated With Specific Methylation Changes: Implication for Molecular Epigenetic Diagnosis. Front Genet **10**, 508.

23. Jayakodi, M., Padmarasu, S., Haberer, G., Bonthala, V.S., Gundlach, H., Monat, C., Lux, T., Kamal, N., Lang, D., Himmelbach, A., Ens, J., Zhang, X.-Q., Angessa, T.T., Zhou, G., Tan, C., Hill, C., Wang, P., Schreiber, M., Boston, L.B., Plott, C., Jenkins, J., Guo, Y., Fiebig, A., Budak, H., Xu, D., Zhang, J., Wang, C., Grimwood, J., Schmutz, J., Guo, G., Zhang, G., Mochida, K., Hirayama, T., Sato, K., Chalmers, K.J., Langridge, P., Waugh, R., Pozniak, C.J., Scholz, U., Mayer, K.F.X., Spannagl, M., Li, C., Mascher, M., and Stein, N. (2020). The barley pan- genome reveals the hidden legacy of mutation breeding. Nature **588**, 284–289.

24. Kanehisa, M., Araki, M., Goto, S., Hattori, M., Hirakawa, M., Itoh, M., Katayama, T., Kawashima, S., Okuda, S., Tokimatsu, T., and Yamanishi, Y. (2008). KEGG for linking genomes to life and the environment. Nucleic Acids Res **36**, D480–484.

25. Katzen, F. (2007). Gateway((R)) recombinational cloning: a biological operating system. Expert Opin Drug Discov **2**, 571–589.

26. Kim, D., Langmead, B., and Salzberg, S.L. (2015). HISAT: a fast spliced aligner with low memory requirements. Nature Methods **12**, 357–360.

27. Kitada, T., Kuryan, B.G., Tran, N.N., Song, C., Xue, Y., Carey, M., and Grunstein, M. (2012). Mechanism for epigenetic variegation of gene expression at yeast telomeric heterochromatin. Genes Dev **26**, 2443–2455.

28. Krueger, F., and Andrews, S.R. (2011). Bismark: a flexible aligner and methylation caller for Bisulfite-Seq applications. Bioinformatics **27**, 1571–1572.

29. Kuo, Y.-T., Câmara, A.S., Schubert, V., Neumann, P., Macas, J., Melzer, M., Chen, J., Fuchs, J., Abel, S., Klocke, E., Huettel, B., Himmelbach, A., Demidov, D., Dunemann, F., Mascher, M., Ishii, T., Marques, A., and Houben, A. (2023). Holocentromeres can consist of merely a few megabase-sized satellite arrays. Nature Communications **14**, 3502.

30. Langmead, B., and Salzberg, S.L. (2012). Fast gapped-read alignment with Bowtie 2. Nature Methods **9**, 357–359.

31. Lavington, E., and Kern, A.D. (2017). The Effect of Common Inversion Polymorphisms In(2L)t and In(3R)Mo on Patterns of Transcriptional Variation in Drosophila melanogaster. G3 (Bethesda) **7**, 3659–3668.

32. Lopez-Delisle, L., Rabbani, L., Wolff, J., Bhardwaj, V., Backofen, R., Grüning, B., Ramírez, F., and Manke, T. (2020). pyGenomeTracks: reproducible plots for multivariate genomic datasets Bioinformatics **37**, 422–423.

33. Lopez, F.B., Fort, A., Tadini, L., Probst, A.V., McHale, M., Friel, J., Ryder, P., Pontvianne, F.D.R., Pesaresi, P., Sulpice, R., McKeown, P., Brychkova, G., and Spillane, C. (2021). Gene dosage compensation of rRNA transcript levels in Arabidopsis thaliana lines with reduced ribosomal gene copy number. Plant Cell **33**, 1135–1150.

34. Love, M.I., Huber, W., and Anders, S. (2014). Moderated estimation of fold change and dispersion for RNA-seq data with DESeq2. Genome Biology **15**, 550.

35. Mandáková, T., and Lysak, M.A. (2016). Chromosome Preparation for Cytogenetic Analyses in Arabidopsis. Curr Protoc Plant Biol **1**, 43–51.

36. Martinez-Zapater, J.M., Estelle, M.A., and Somerville, C.R. (1986). A highly repeated DNA sequence in Arabidopsis thaliana. Molecular and General Genetics MGG **204**, 417–423.

37. Mohannath, G., Pontvianne, F., and Pikaard, C.S. (2016). Selective nucleolus organizer inactivation in *Arabidopsis* is a chromosome position-effect phenomenon. Proceedings of the National Academy of Sciences **113**, 13426–13431.

38. Naish, M., Alonge, M., Wlodzimierz, P., Tock, A.J., Abramson, B.W., Schmücker, A., Mandáková, T., Jamge, B., Lambing, C., Kuo, P., Yelina, N., Hartwick, N., Colt, K., Smith, L.M., Ton, J., Kakutani, T., Martienssen, R.A., Schneeberger, K., Lysak, M.A., Berger, F., Bousios, A., Michael, T.P., Schatz, M.C., and Henderson, I.R. (2021). The genetic and epigenetic landscape of the *Arabidopsis* centromeres. Science **374**, eabi7489.

39. Naseeb, S., Carter, Z., Minnis, D., Donaldson, I., Zeef, L., and Delneri, D. (2016). Widespread Impact of Chromosomal Inversions on Gene Expression Uncovers Robustness via Phenotypic Buffering. Molecular Biology and Evolution **33**, 1679–1696.

40. Puchta, H., and Houben, A. (2024). Plant chromosome engineering – past, present and future. New Phytologist **241**, 541–552.

41. Ramírez, F., Ryan, D.P., Grüning, B., Bhardwaj, V., Kilpert, F., Richter, A.S., Heyne, S., Dündar, F., and Manke, T. (2016). deepTools2: a next generation web server for deep-sequencing data analysis. Nucleic Acids Research **44**, W160–W165.

42. Robinson, J.T., Thorvaldsdóttir, H., Winckler, W., Guttman, M., Lander, E.S., Getz, G., and Mesirov, J.P. (2011). Integrative genomics viewer. Nat Biotechnol **29**, 24–26.

43. Rönspies, M., Dorn, A., Schindele, P., and Puchta, H. (2021). CRISPR–Cas-mediated chromosome engineering for crop improvement and synthetic biology. Nature Plants **7**, 566–573.

44. Rönspies, M., Schindele, P., Wetzel, R., and Puchta, H. (2022a). CRISPR-Cas9-mediated chromosome engineering in Arabidopsis thaliana. Nat Protoc **17**, 1332–1358.

45. Rönspies, M., Schmidt, C., Schindele, P., Lieberman-Lazarovich, M., Houben, A., and Puchta, H. (2022b). Massive crossover suppression by CRISPR-Cas-mediated plant chromosome engineering. Nat Plants **8**, 1153–1159.

46. Schmidt, C., Fransz, P., Rönspies, M., Dreissig, S., Fuchs, J., Heckmann, S., Houben, A., and Puchta, H. (2020). Changing local recombination patterns in Arabidopsis by CRISPR/Cas mediated chromosome engineering. Nature Communications **11**, 4418.

47. Schwartz, C., Lenderts, B., Feigenbutz, L., Barone, P., Llaca, V., Fengler, K., and Svitashev, S. (2020). CRISPR–Cas9-mediated 75.5-Mb inversion in maize. Nature Plants **6**, 1427–1431.

48. Sherman, B.T., Hao, M., Qiu, J., Jiao, X., Baseler, M.W., Lane, H.C., Imamichi, T., and Chang, W. (2022). DAVID: a web server for functional enrichment analysis and functional annotation of gene lists (2021 update). Nucleic Acids Res **50**, W216–w221.

49. Stark, R., and Brown, G.D. (2012). DiffBind : Differential binding analysis of ChIP-Seq peak data.

50. Steinert, J., Schiml, S., Fauser, F., and Puchta, H. (2015). Highly efficient heritable plant genome engineering using Cas9 orthologues from Streptococcus thermophilus and Staphylococcus aureus. The Plant Journal **84**, 1295–1305.

51. Tang, D., Chen, M., Huang, X., Zhang, G., Zeng, L., Zhang, G., Wu, S., and Wang, Y. (2023). SRplot: A free online platform for data visualization and graphing. PLoS One **18**, e0294236.

52. Termolino, P., Falque, M., Aiese Cigliano, R., Cremona, G., Paparo, R., Ederveen, A., Martin, O.C., Consiglio, F.M., and Conicella, C. (2019). Recombination suppression in heterozygotes for a pericentric inversion induces the interchromosomal effect on crossovers in Arabidopsis. Plant J **100**, 1163–1175.

53. Wang, Y., Qu, Z., Fang, Y., Chen, Y., Peng, J., Song, J., Li, J., Shi, J., Zhou, J.-Q., and Zhao, Y. (2023). Chromosome territory reorganization through artificial chromosome fusion is compatible with cell fate determination and mouse development. Cell Discovery **9**, 11.

54. Weiss, T., Crisp, P.A., Rai, K.M., Song, M., Springer, N.M., and Zhang, F. (2022). Epigenetic features drastically impact CRISPR–Cas9 efficacy in plants. Plant Physiology **190**, 1153–1164.

55. Zhang, Y., Liu, T., Meyer, C.A., Eeckhoute, J., Johnson, D.S., Bernstein, B.E., Nusbaum, C., Myers, R.M., Brown, M., Li, W., and Liu, X.S. (2008). Model-based Analysis of ChIP-Seq (MACS). Genome Biology **9**, R137.

56. Zhou, Y., Yu, Z., Chebotarov, D., Chougule, K., Lu, Z., Rivera, L.F., Kathiresan, N., Al-Bader, N., Mohammed, N., Alsantely, A., Mussurova, S., Santos, J., Thimma, M., Troukhan, M., Fornasiero, A., Green, C.D., Copetti, D., Kudrna, D., Llaca, V., Lorieux, M., Zuccolo, A., Ware, D., McNally, K., Zhang, J., and Wing, R.A. (2023). Pan-genome inversion index reveals evolutionary insights into the subpopulation structure of Asian rice. Nat Commun **14**, 1567.

